## Supplemental Information for "Stage-specific expression patterns and co-targeting relationships among miRNAs in the developing mouse cerebral cortex"

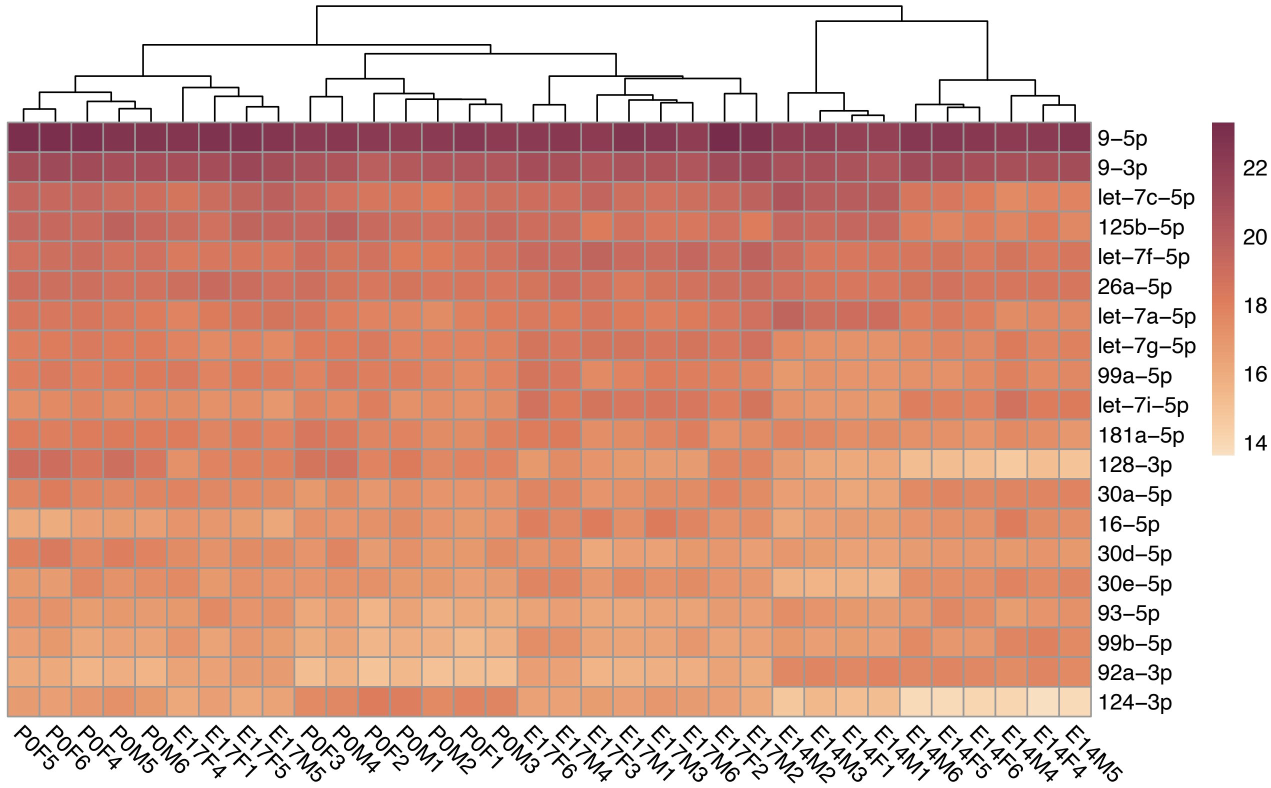


**Fig S1.** Top 20 most highly expressed miRNAs in E14, E17 and P0 mouse cortical samples. Values in the heatmap correspond to log2-transformed normalized counts. M – males, F – females.

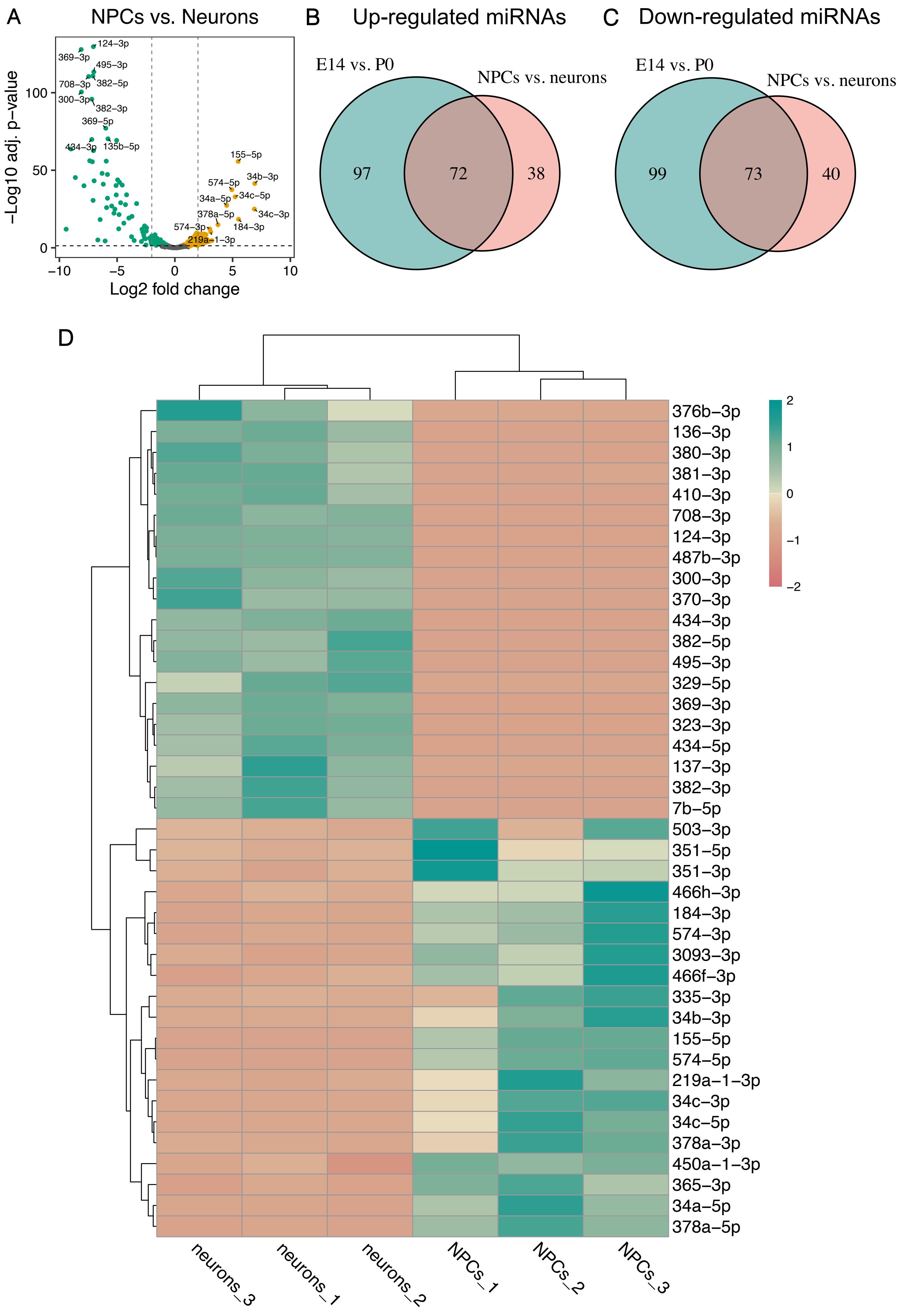
A
NPCs vs. Neurons
−Log10 adj. p−value
Log2 fold change
124−3p
369−3p
495−3p
708−3p
382−5p
300−3p
382−3p
369−5p
434−3p
135b−5p
155−5p
574−5p
34b−3p
34a−5p
34c−5p
378a−5p
34c−3p
574−3p
184−3p
219a−1−3p
B
Up-regulated miRNAs
E14 vs. P0
NPCs vs. neurons
97
72
38
C
Down-regulated miRNAs
E14 vs. P0
NPCs vs. neurons
99
73
40
D
376b−3p
136−3p
380−3p
381−3p
410−3p
708−3p
124−3p
487b−3p
300−3p
370−3p
434−3p
382−5p
495−3p
329−5p
369−3p
323−3p
434−5p
137−3p
382−3p
7b−5p
503−3p
351−5p
351−3p
466h−3p
184−3p
574−3p
3093−3p
466f−3p
335−3p
34b−3p
155−5p
574−5p
219a−1−3p
34c−3p
34c−5p
378a−3p
450a−1−3p
365−3p
34a−5p
378a−5p
neurons_3
neurons_1
neurons_2
NPCs_1
NPCs_2
NPCs_3

**Fig S2.** Differential expression analysis of miRNAs in neuronal progenitor cells (NPCs) vs. neurons. **A** Volcano plot of differentially expressed miRNAs. Orange dots correspond to miRNAs up-regulated in NPCs, blue dots signify miRNAs that are up-regulated in neurons. Non-significant results are shown in gray. **B-C** Venn diagrams showing the overlap of miRNAs that were up- or down-regulated in bulk cortical samples from E14 vs. P0 and NPCs vs. neurons. **D** Heatmap showing the expression levels of the top 20 down- and top 20 up-regulated miRNAs in NPCs vs. neurons. Values correspond to z-scores of normalized miRNA expression.

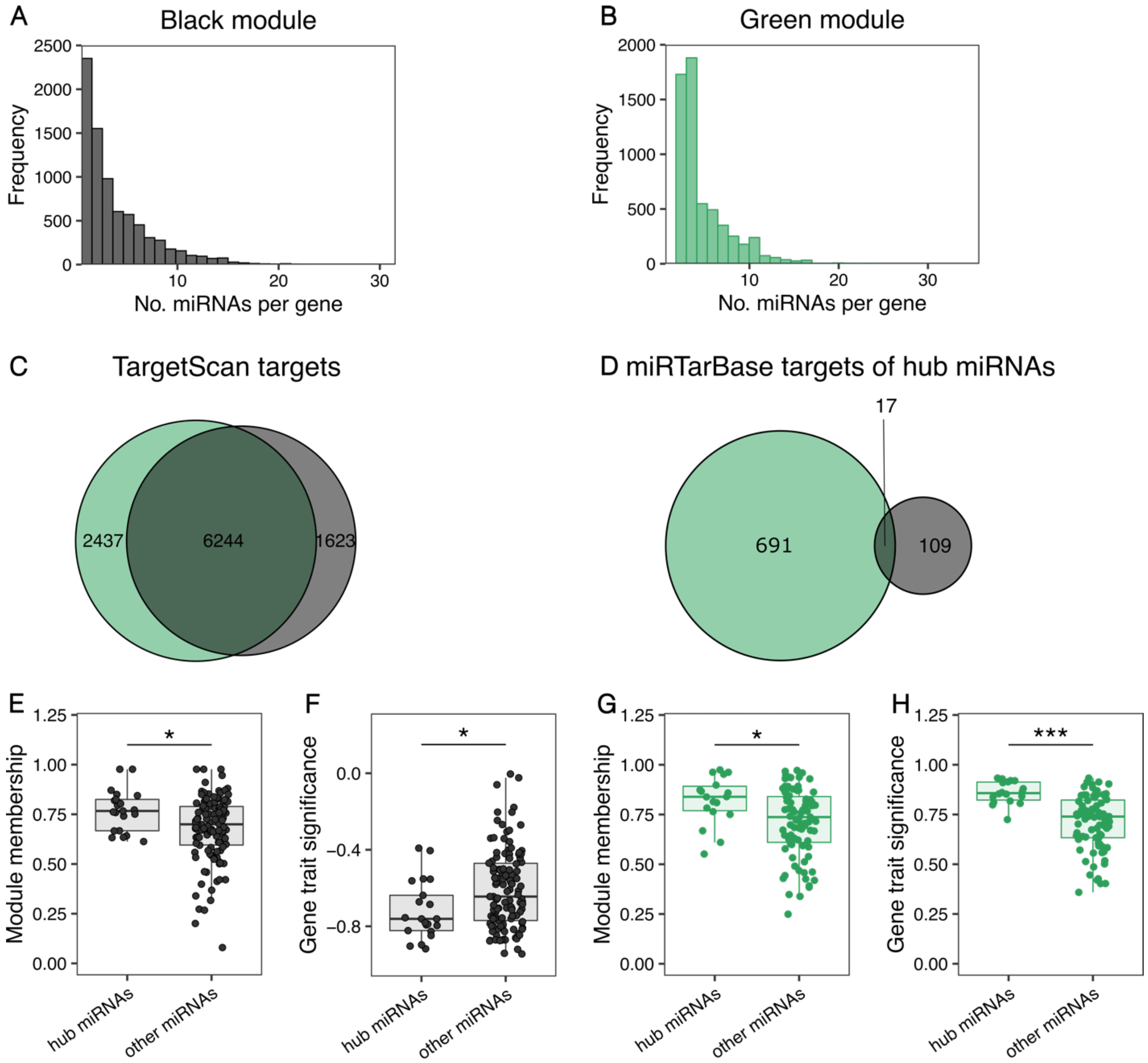


**Fig S3.** Histograms show the frequency distribution for the number of miRNAs targeting a gene for the black (**A)** and green (**B**) modules. **C** Venn diagram with the overlap of common genes targeted by miRNAs in the black and green modules. Target predictions were obtained from TargetScan using conserved miRNA families and binding sites. **D** Venn diagram with validated targets for conserved hub miRNAs from the black and green module obtained from miRTarBase. **E-H** Box plots with the module membership and gene-trait significance values for the hub miRNAs and the remaining miRNAs in the black and green modules. ***$p<0.001$, * $p<0.05$, unpaired t-test.

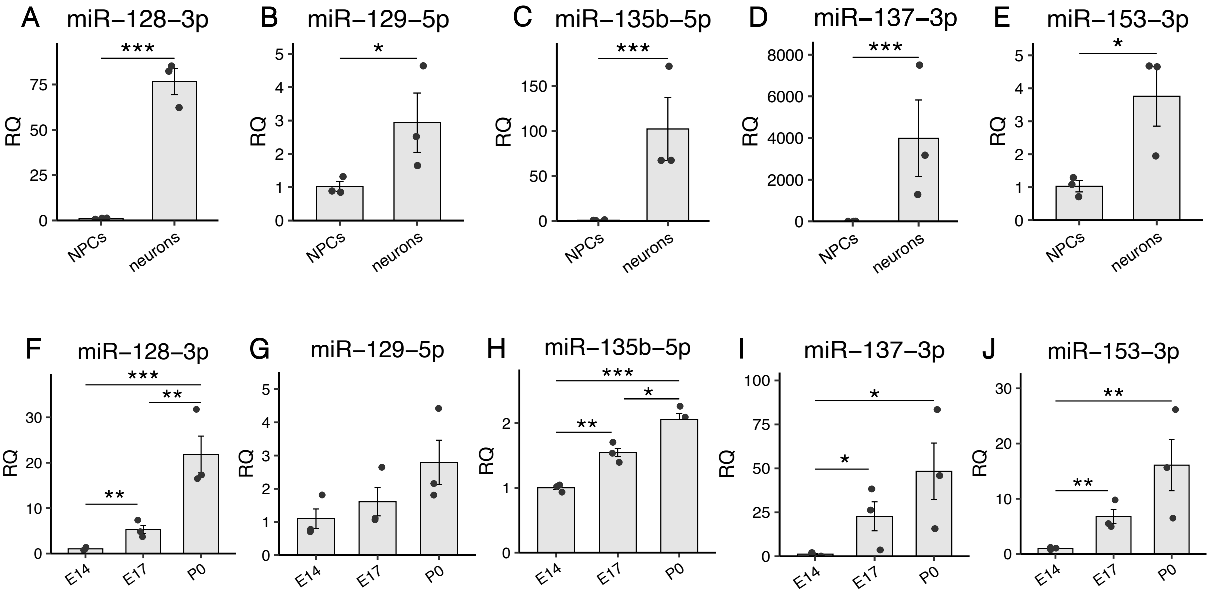


**Fig S4. Relative quantification of miRNA expression using RT-qPCR**

**A-E** Expression levels of miRNAs selected for luciferase experiments in neuronal progenitor cells (NPCs) versus neurons. Relative quantification (RQ) values were normalized to the mean expression in NPCs. **F-J** Expression levels in cortical samples at E14, E17 and P0. RQ values were normalized to the mean expression in the E14 group.

Data are shown as mean ± standard error of the mean and individual values. *p<0.05, ** p<0.01, *** p<0.001, unpaired t-test of log-transformed RQ values.

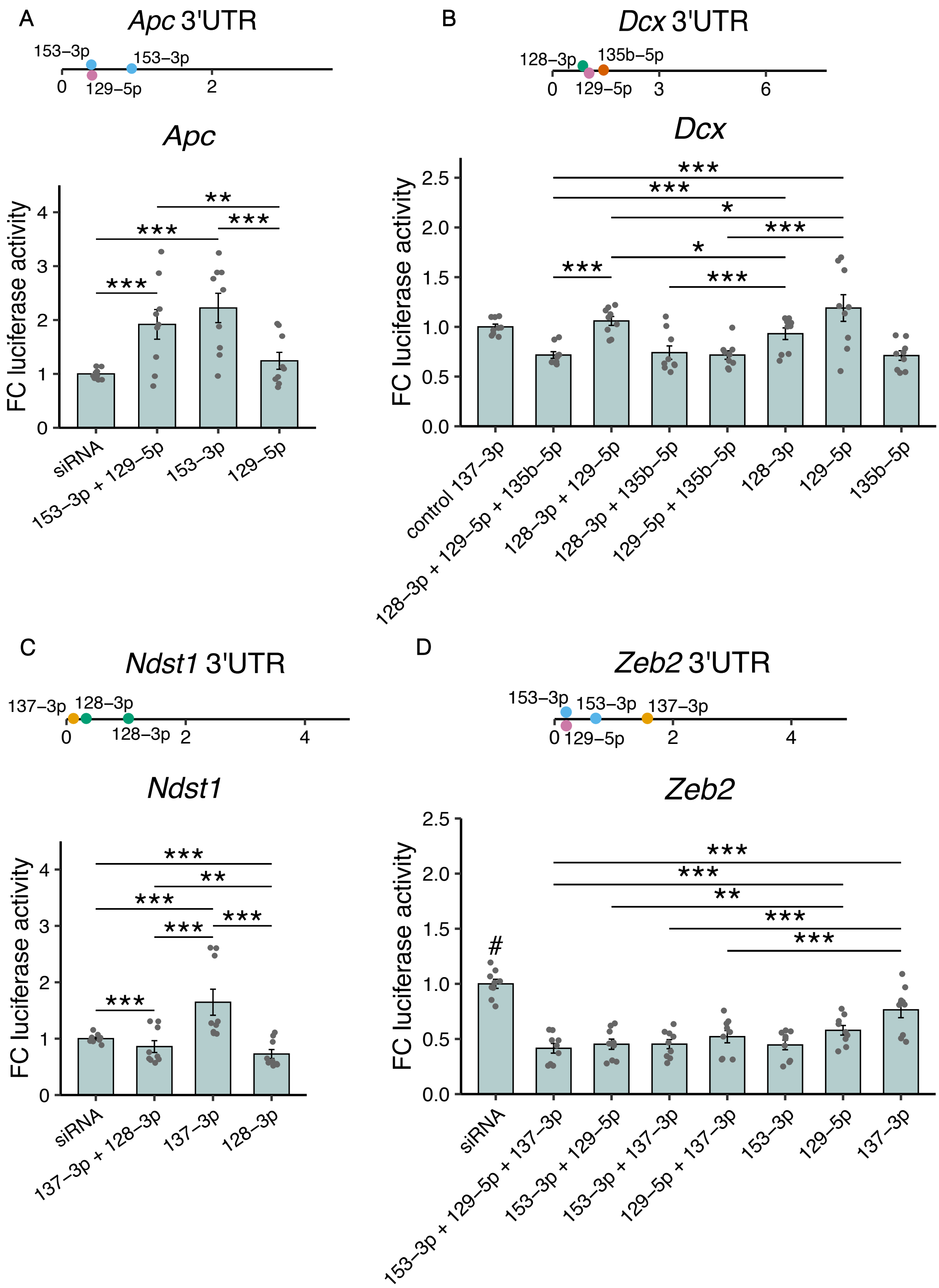


**Fig S5. Luciferase activity in lysates of HEK293 cells transfected with plasmids containing 3' UTR fragments of target genes.** Lysates were co-transfected with different combinations of miRNA mimics. Fold change (FC) of luciferase activity was obtained by calculating the ratio of *Renilla* luciferace and firefly luciferase activity and then normalizing to the mean of the control siRNA or miRNA group. Locations of the binding sites in the 3' UTR of the respective gene are represented by colored dots. The length of the 3' UTRs is indicated in kbp. Data are shown as mean ± standard error of the mean and individual values. *p<0.05, ** p<0.01, *** p<0.001, only statistical comparisons containing groups with at least two miRNA mimics are shown, #p<0.05 siRNA control vs. all other groups, two-way analysis of variance followed by Tukey's post-hoc test.

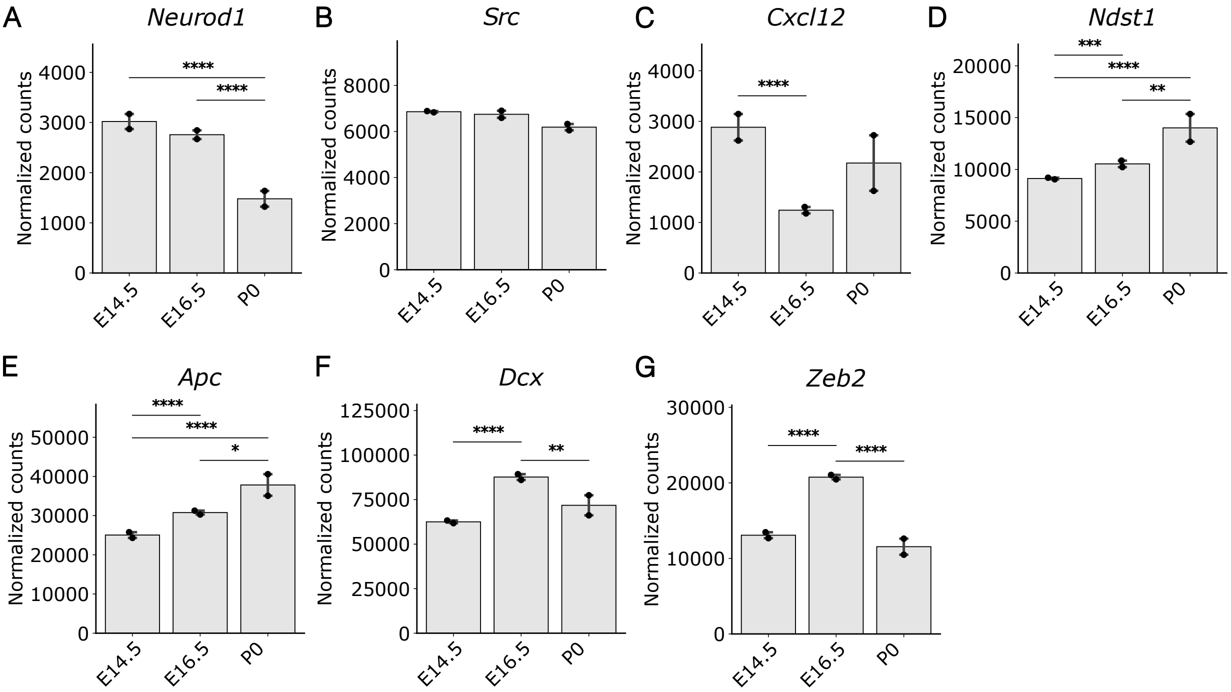


**Fig S6. Expression of target genes from the luciferase assay at different developmental time points.** Bar plots show DESeq2-normalized counts for target genes that were used in the luciferase experiments at E14.5, E16.5 and P0 of cortical development. Data are depicted as mean ± standard error of the mean ** $p<0.01$, ***$p<0.001$, ***$p<0.0001$, Wald test from DESeq2. The expression values of genes were obtained by reprocessing data from the study by Weyn-Vanhentenryck et al. [1]

**Table S1. Primers used for PCR and nested PCR of target genes used in luciferase assays.**

| Primer name | Sequence 5' to 3' | PCR product size |
|---|---|---|
| **Neurod1_XhoI_forw** | ACGTCTCGAGGCCTTTGGAAGAAACAGGGG | 274 bp |
| **Neurod1_NotI_rev** | ACGTGCGGCCGCGGTCACAGGTAGTAAAATGCTGG | |
| **Neurod1_lo_XhoI_for** | ACGTCTCGAGCGTCAGTTTCACTATTCCCGG | 1188 bp |
| **Neurod1_lo_NotI_rev** | ACGTGCGGCCGCCAGCACTTATTCTGGACTGCA | |
| **Zeb2_XhoI_forw** | ACGTCTCGAGCCAGGAAGCTGTAGAGAGGG | 1555 bp |
| **Zeb2_NotI_rev** | ACGTGCGGCCGCCAGGATCAGTTGAGAAAAGCTGT | |
| **Dcx_XhoI_forw** | ACGTCTCGAGGTTTGGGGTACATGATGTCACA | 1022 bp |
| **Dcx_NotI_rev** | ACGTGCGGCCGCCATCAGCAATGCCACCAAGT | |
| **Ndst1_XhoI_forw** | ACGTCTCGAGCTTGTGTTCGCAGGGATGTC | 1221 bp |
| **Ndst1_NotI_rev** | ACGTGCGGCCGCAGGACCCTTCAAGACTTCGC | |
| **Src_XhoI_forw** | ACGTCTCGAGCCCTGTGTGTGTGTGTTTGT | 484 bp |
| **Src_NotI_rev** | ACGTGCGGCCGCTACAACAAGTCTGGGTCCCC | |
| **Cxcl12_XhoI_forw** | ACGTCTCGAGACAGTGGGGATTCTGGGTTC | 407 bp |
| **Cxcl12_NotI_rev** | ACGTGCGGCCGCACGGTAGGAGGTTTACAGCA | |
| **Nipbl_XhoI_forw** | ACGTCTCGAGACATGCAGCCAAATTTACAGG | 748 bp |
| **Nipbl_NotI_rev** | ACGTGCGGCCGCAAGTCAGCCTGTACAAACTGT | |
| **Cited2_XhoI_forw** | ACGTCTCGAGCACAAACTGCCATCTCGCTT | 311 bp |
| **Cited2_NotI_rev** | ACGTGCGGCCGCCCTAAAAAGCTTTCAACACAGTAG | |
| **Nfib_1_XhoI_forw** | ACGTCTCGAGACATTACGTGCCTTGCCTTG | 2529 bp |
| **Nfib_1_NotI_rev** | ACGTGCGGCCGCCTTCTCTCTCCTCGCAGCTT | |
| **Nfib_2_XhoI_forw** | ACGTCTCGAGATGCATTCTTCATCGAGGGC | 957 bp |
| **Nfib_2_NotI_rev** | ACGTGCGGCCGCGTCTGTAGCATAGCTCATTT | |
| **Apc_nested_forw** | GAGCCCAAAGTCCTAAACGC | 1304 bp |

| | | |
|---|---|---|
| **Apc_nested_rev** | CGGAGAAGATGACGGGGTAA | |
| **Apc_XhoI_forw** | ACGTCTCGAGCTGGTCATTTTGGGAGGCAC | 962 bp |
| **Apc_XhoI_forw** | ACGTGCGGCCGCGCCCTTATGTCCAGTGCCTA | |

**Table S2. Primers used for targeted in vitro mutagenesis of miRNA binding sites.**

| Primer name | Sequence 5' to 3' |
|---|---|
| **Neurod1_Mut137_forw** | CTGATCGGGATAAAAAAAATCACAAACGATAATTAGGATC |
| **Neurod1_Mut137_rev** | GATCCTAATTATCGTTTGTGATTTTTTTTATCCCGATCAG |
| **Neurod1_Mut153_forw** | TAATTAGGATCTGTACAATTTTTAAACTAGTAATGGGCC |
| **Neurod1_Mut153_rev** | GGCCCATTACTAGTTTAAAAATTGTACAGATCCTAATTA |
| **Cxcl12_Mut135b_forw** | ATATATTTGAAGTGGAGCTACAGTAATGCCAGTAGAT |
| **Cxcl12_Mut135b_rev** | ATCTACTGGCATTACTGTAGCTCCACTTCAAATATAT |
| **Cxcl12_Mut137_forw** | CTGTGACATTATATGCACTAACGATAAAATGCTAATTGTTTC |
| **Cxcl12_Mut137_rev** | GAAACAATTAGCATTTTATCGTTAGTGCATATAATGTCACAG |
| **Src_Mut137_forw** | CCATTGCCCCATCACAACGATAATGTCCCCGCTACTGG |
| **Src_Mut137_rev** | CCAGTAGCGGGGACATTATCGTTGTGATGGGGCAATGG |
| **Src_Mut153_forw** | GTAGATTTCAGATGACTGATCAGAGGCCTTGGGGACC |
| **Src_Mut153_rev** | GGTCCCCAAGGCCTCTGATCAGTCATCTGAAATCTAC |

**References**

1. Weyn-Vanhentenryck SM, Feng H, Ustianenko D, Duffié R, Yan Q, Jacko M, Martinez JC, Goodwin M, Zhang X, Hengst U, et al: **Precise temporal regulation of alternative splicing during neural development.** *Nature Communications* 2018, **9:**2189.
